# Uncovering functional connectivity patterns predictive of cognition in youth using interpretable predictive modeling

**DOI:** 10.1101/2025.03.09.642155

**Authors:** Hongming Li, Matthew Cieslak, Taylor Salo, Russell T. Shinohara, Desmond J. Oathes, Christos Davatzikos, Theodore D. Satterthwaite, Yong Fan

## Abstract

Functional MRI studies have identified functional connectivity (FC) patterns associated with behavioral traits using whole-brain or region-wise predictive models. However, whole-brain approaches often suffer from limited generalizability and interpretability due to the high-dimensionality of FC data. Conversely, region-wise models inherently isolate predictions, ineffective for characterizing contributions of the whole brain FC patterns in predicting a target trait. In this study, we propose an interpretable predictive model that learns fine-grained FC patterns predictive of behavioral traits, jointly at the regional and participant levels, to characterize the overall association of FC patterns with a target trait. Our model learns both a relevance score and a dedicated prediction model for each brain region, then integrates the regional predictions to generate a participant-level prediction, capturing the collective association of FC patterns with the trait. We validated our method using FC data from 6798 participants in the Adolescent Brain and Cognitive Development (ABCD) study for predicting cognition. Our interpretable predictive model identified the cingulo-parietal, retrosplenial-temporal, dorsal attention, salience, and cingulo-opercular networks as collectively predictive of cognitive traits. The interpretable model significantly improved prediction accuracy and facilitated the characterization of fine-grained differences in FC patterns across cognitive domains. Furthermore, the learned relevance scores enhanced region-wise predictions of longitudinal cognitive measures in the ABCD cohort and cognitive traits in an external Human Connectome Project Development (HCP-D) cohort. These findings suggest that our method effectively characterizes generalizable and fine-grained FC patterns linked to cognition in youth.

## Introduction

Neurocognitive development is a protracted process, unfolding alongside the maturation of brain functional organization during childhood and adolescence [1, 2]. Individual differences in youth neurocognition are linked to critical outcomes such as academic performance and quality of life [2], while neurocognitive deficits are associated with trans-diagnostic mental illness [3]. As such, it is crucial to deepen our understanding of the relationship between individual cognitive differences and brain functional organization in youth.

Functional magnetic resonance imaging (fMRI) studies have used machine learning (ML)-based predictive modeling to identify functional network patterns associated with behavioral traits. Global models, which use whole-brain functional connectivity (FC) measures as input features, are common. Inter-individual FC differences have been demonstrated to predict maturity [4], cognition [5-11], personality [7, 11-14], mental health [7, 11, 15-17], as well as sex and gender [18, 19]. While global models can potentially capture interplays across multiple brain regions and their associations with behavior, they face challenges in interpretability and generalizability due to the high dimensionality of whole-brain FC data.

Specifically, global models struggle to capture fine-grained FC information, often relying on post-hoc analyses to identify important brain regions [20]. Common approaches include feature weight-based metrics, which define regional importance as the sum of feature weights from all FCs connected to the region [7, 18], and sensitivity analysis, which assesses changes in prediction performance after excluding specific regions [21, 22]. While these post-hoc techniques provide helpful insights, they may not accurately reflect the model’s decision-making process, limiting their suitability for high-stakes biological and clinical applications [23].

In contrast, region-wise models offer some inherent interpretability, with prediction performance directly highlighting significant regions [20]. For example, regions with higher prediction accuracy are considered more strongly linked with the traits of interest. However, these models inherently ignore the influence of multiple-region interactions on behavior, as prediction is performed separately for each region. Such models have been used to explore associations between FC and various behaviors/phenotypes, including sex differences and borderline personality traits [12, 24].

To bridge the gap between global and region-wise approaches for identifying the brain-behavior associations, we propose an interpretable predictive modeling method that learns fine-grained FC patterns predictive of behavioral traits at the participant level. This approach captures both regional and global FC associations with the target trait of interest, enhancing interpretability and improving prediction accuracy. Specifically, our model learns a relevance score and a dedicated prediction model for each brain region, then integrates these regional predictions to generate a participant-level prediction. The model is optimized end-to-end by collaboratively minimizing the differences between the predicted and measured traits at both the regional and participant levels. We validated our method using FC data from 6798 participants in the Adolescent Brain and Cognitive Development (ABCD) study for predicting cognition. Results demonstrate that brain regions within the cingulo-parietal, retrosplenial-temporal, dorsal attention, salience, and cingulo-opercular networks are collectively associated with cognitive traits. Our model achieved significantly higher prediction accuracy compared to alternative whole-brain FC-based prediction model and ensembles of region-wise models. Furthermore, the learned relevance scores improved predictions of longitudinal cognitive measures in the ABCD cohort and cognitive traits in an external dataset of 454 participants from the Human Connectome Project Development (HCP-D) cohort. These findings suggest that our method effectively characterizes generalizable and fine-grained FC patterns associated with cognition in youth.

## Methods

### Material

Neuroimaging data and cognitive measures used in this study were drawn from two cohorts: the Adolescent Brain and Cognitive Development study (ABCD; *n*=6798) and the Human Connectome Project Development (HCP-D; *n*=454).

The ABCD study [25] includes 11,878 participants aged 9-10 years at baseline, with fMRI and behavioral data collected across 21 sites in the U.S. Data preprocessed by the ABCD BIDS Community Collection (ABCC, ABCD-3165) [26] were used. After excluding participants with incomplete data or excessive head motion, the final sample included 6798 participants (3367 males and 3431 females).

The HCP-D [27] includes 652 participants recruited across 4 sites in the U.S. to study healthy brain development in children and adolescents. Following previously described inclusion criteria [28], a sample of 454 participants (215 males and 239 females) aged 8-22 years with complete demographic, cognitive assessment, and neuroimaging data was included in this study. Neuroimaging acquisition and preprocessing procedures have been previously described [28].

For both samples, preprocessed fMRI time series were parcellated into 333 cortical regions using the Gordon atlas [29]. For each participant, time series from all resting-state fMRI scans were concatenated to compute FC measures. Pearson’s correlation coefficient between the time series of two cortical regions served as their FC measure.

For the ABCD sample, three factor-analysis based composite scores, including general cognition, executive function, and learning/memory, were used as the baseline cognitive scores [30]. Cognitive scores assessed by a battery of cognitive assessments at a 2-year follow-up were used as longitudinal cognitive measures, including the Picture Vocabulary (PicVocab), Flanker Test (Flanker), Pattern Comparison Processing Speed Task (Pattern), Picture Sequence Memory Task (Picture), Oral Reading Test (Reading), Little Man Task (LMT), and Rey Auditory Verbal Learning Task (RAVLT) [31, 32]. Longitudinal data were available for 4484 participants (2215 males and 2269 females). For the HCP-D sample, three composite scores including fluid cognition, crystalized cognition, and total cognition from the HCP-D data release were used [27].

### Interpretable predictive modeling

An interpretable predictive modeling framework is developed to learn fine-grained FC patterns predictive of behavioral traits at both regional and participant levels, as illustrated in Fig. 1. This framework constructs prediction models based on regional FC measures and integrates these region-wise models in an end-to-end learning process, effectively characterizing the contributions of both regional and global FCs to the target trait.

**Fig. 1.**
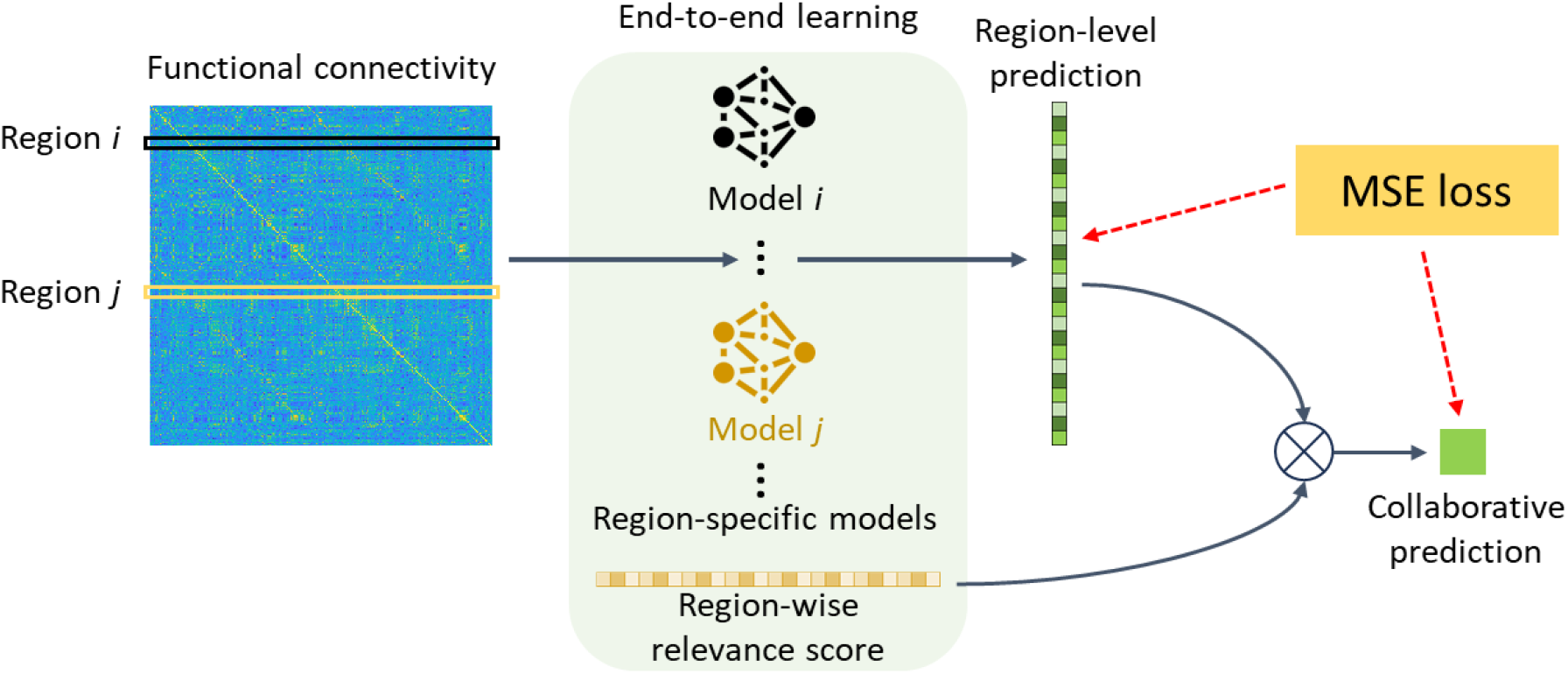
Interpretable predictive modeling of FC-behavior association. Our model learns fine-grained FC patterns predictive of behavioral traits at both regional and participant levels, capturing the overall FC-behavior association. Region-wise predictions are integrated using region-wise relevance scores, yielding a participant level prediction and an interpretable measure of each region’s contribution. These regional predictions and relevance scores are optimized collaboratively using a mean square error (MSE) based loss to enhance prediction performance.

Given *N* participants, each with a matrix *C*^*i*^ ∈ *R*^*M*×*M*^, *i* = 1,2, …, *N* for quantifying FC measures across *M* brain regions, we denote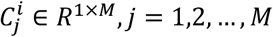, the *j*-th row of *C*^*i*^, as the FC profile of the *j*-th brain region for participant *i*. This profile refers to the FC measures between the *j*-th region and all other regions. A region-specific prediction model *f*_*j*_ is used to map the FC profile of the *j*-th region to the behavioral trait of interest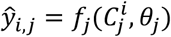, where *θ*_*j*_ represents the model parameters for the *j*-th region to be optimized, and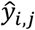 represents the predicted trait for the *j*-th region of participant *i*. Participant-level prediction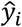is obtained by integrating the region-level predictions:

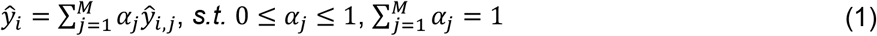

where *α*_*j*_ is the relevance score for the *j*-th region, indicating its contribution to the overall prediction. The regional relevance scores {*α*_*j*_}_*j*=1,2,…,*M*_, along with the regional model {*f*_*j*_}_*j*=1,2,…,*M*_, are learned jointly during model training.

To capture FC-trait relationships at both regional and global levels, the model parameters {*θ*_*j*_}_*j*=1,…,*M*_ and relevance scores {*α*_*j*_}_*j*=1,…,*M*_ are optimized jointly by minimizing the differences between the predicted and measured traits:

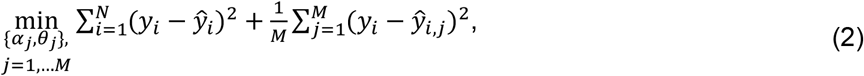

where *y*_*i*_ refers to the measured trait for participant *i*, the first term is the mean square error (MSE) loss at the participant level, and the second term is the averaged MSE loss at the region level.

The proposed model was constructed and optimized under a neural network setting using PyTorch [33]. The region-specific prediction model *f*_*j*_ was implemented using a fully connected layer, with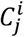as input and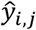as output for participant *i*. The relevance scores {*α*_*j*_}_*j*=1,…,*M*_ were implemented as trainable network parameters and was followed by a normalization step of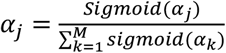 The initial value of {*α*_*j*_}_*j*=1,…,*M*_ was set to 0 so that equal relevance score (^1^ after normalization) was assigned to each brain region at the beginning of model training. For the model training, the Adam optimizer [34] was adopted to optimize all the network parameters with a learning rate of 0.0005, the batch size was set to 32 and the number of epochs set to 200.

### Model training and evaluation

Model training and evaluation were conducted using five-fold stratified cross-validation to mitigate the effects of confounding factors on the prediction. Specifically, stratified data splitting was utilized to obtain the training and testing data splits with confounding factors controlled, so that distributions of the confounding factors in training and testing data splits were matched. For the ABCD cohort, these factors included age, sex, race, and site information, while for the HCP-D cohort they included age and sex.

FC measures derived from baseline fMRI scans were used as features, while baseline and follow-up cognitive measures were used as target variables for prediction. Prediction accuracy was assessed using the mean absolute error (MAE) and partial correlation coefficients between predicted and measured cognitive measures. The partial correlation coefficients were obtained by controlling for age, sex, and head motion.

Our interpretable predictive model (Region-int) was compared against two common fMRI-based baselines: 1) ridge regression on whole-brain FC measures (Whole-FC), and 2) averaged ridge regression models for each brain region (Region-avg). Particularly, the upper-triangle of the FC matrix was flattened and used as whole-brain FC measures to train the Whole-FC model. The Region-avg model was implemented by first training a dedicated ridge regression model for each brain region with its FC profile as features and then averaging the region-wise predictions to obtain the final prediction for the testing participants. All models underwent five-fold cross-validation with identical training/testing data splits. In all ridge regression models, the hyperparameter *λ* was optimized using nested cross-validation within the training dataset.

The ABCD cohort served as the primary dataset for evaluating model performance and the learned region-wise relevance scores. The HCP-D cohort served as an external validation dataset to further evaluate the generalizability of the relevance scores in capturing the associations between regional FC profiles and cognition. The baseline models in ABCD and HCP-D cohorts were trained separately for their specific cognitive measures.

## Results

### Interpretable predictive modeling better predicts cognition

The prediction performance of different models on the ABCD cohort is illustrated in Fig. 2A. The Region-int and the Region-avg model significantly outperformed the Whole-FC model across all three cognitive domains (*p*<0.042, corrected *t*-test). To compare two predictive models, the corrected *t*-test [35, 36] was utilized to compare their accuracy measures across five testing folds. The correlation between measured and predicted values obtained by the proposed Region-int model across testing folds was 0.477 ± 0.015 (mean±std), 0.165 ± 0.016, and 0.262 ± 0.027 for general cognition, executive function, and learning/memory, respectively. These results were better than those obtained by the Region-avg model, which had correlation measures of 0.429 ± 0.021, 0.159 ± 0.027, and 0.243 ± 0.025, particularly for general cognition (*p*<0.001, corrected *t*-test). Consistent comparison results were found regarding the MAE measures. The improved prediction performance suggests that the interpretable predictive modeling can better learn the associations between FC profiles and cognition.

**Fig. 2.**
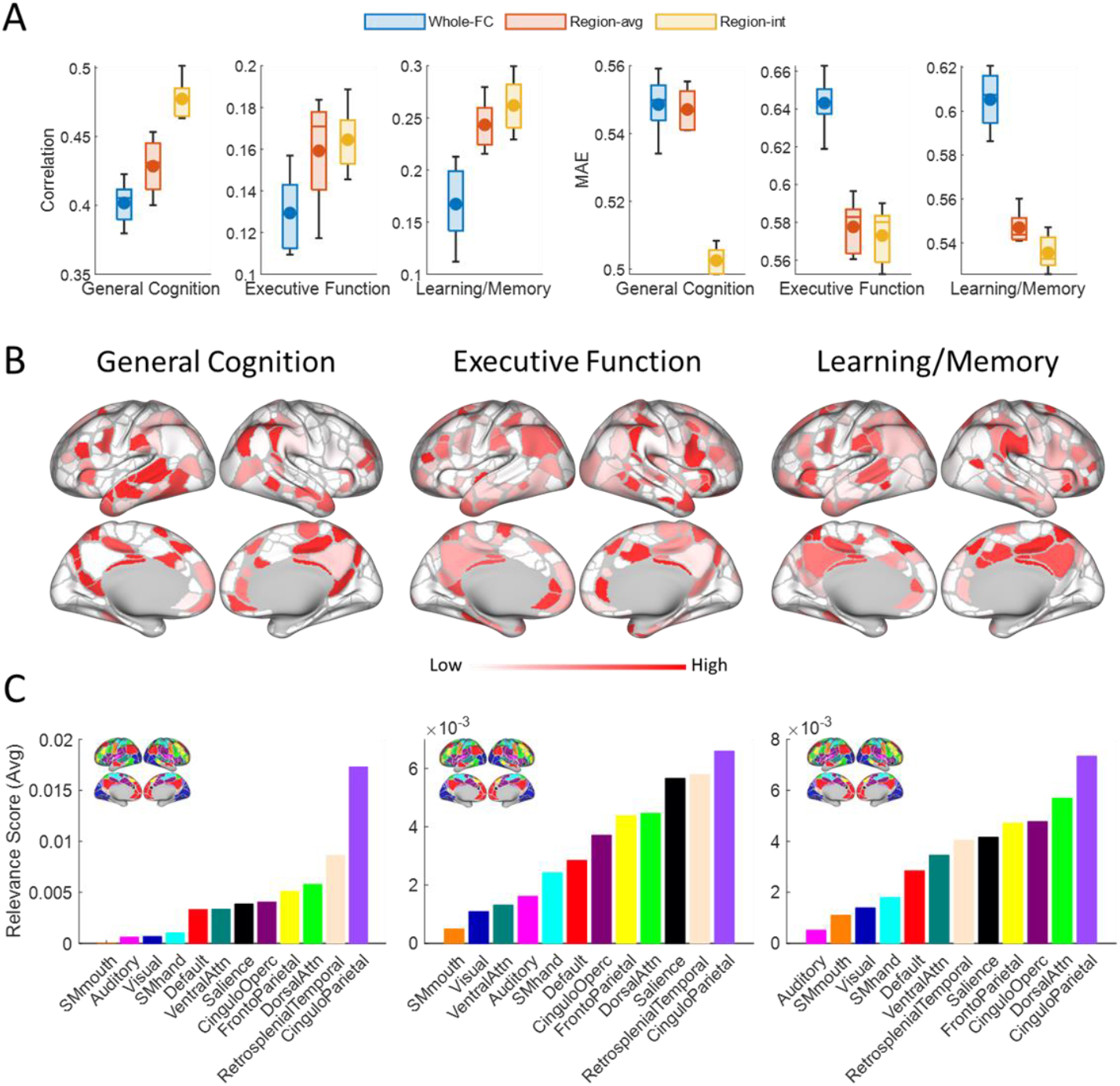
The interpretable predictive modeling captures the FC-cognition associations. (A) The Region-int modeling obtained better prediction accuracy compared to conventional whole-brain FC based prediction and averaged region-wise prediction. The boxplot shows the distribution of prediction accuracy over five-fold cross-validation, with the ‘-’ and ‘•’ markers representing the median and mean, respectively. (B) Relevance scores for different cognitive domains learned by the Region-int model. (C) Distributions of the relevance scores across functional networks for different cognitive domains.

### Fine-grained FC patterns linked to cognition

In addition to improved prediction accuracy, the interpretable predictive model identified fine-grained FC patterns associated with cognition by assigning a relevance score to each cortical region, with higher scores indicating stronger associations between the region’s FC profile and cognition. As shown in Fig. 2B, cortical regions with high relevance to cognition were mainly situated in the association cortex, and the associated FC patterns varied across cognitive domains. To better understand their spatial distribution across the cortex, the region-wise relevance scores were summarized across functional networks (Fig. 2C). On average, regions in the cingulo-parietal network exhibited the highest relevance to all cognitive domains, especially general cognition. The retrosplenial-temporal and dorsal attention networks were also highly associated with general cognition, while the retrosplenial-temporal and salience networks showed higher relevance for executive function, and the dorsal attention and cingulo-opercular networks for learning/memory.

We also found the relevance scores provided additional information about the FC-cognition associations beyond what was revealed by the region-wise prediction accuracy. As shown in Fig. 3A, brain regions within the cingulo-parietal and retrosplenial-temporal networks exhibited high relevance scores across cognitive domains, even though they were not highlighted by the region-wise accuracy, especially for executive function and learning/memory. We further investigated the similarity of FC-cognition association maps quantified by the interpretable predictive model’s relevance scores and the region-wise ridge regression models’ prediction accuracy across cognitive domains. Spearman’s rank correlation was used to measure the similarity between pairs of FC-cognition association maps, with each map thresholded at the 50% percentile to eliminate the influences of brain regions with low relevance or prediction accuracy. While FC-cognition association maps were significantly correlated (*p*<0.0001, Spin test [37, 38]), the relevance scores exhibited greater diversity across cognitive domains, with lower similarity measures of 0.384, 0.634, and 0.423 between general cognition and executive function, general cognition and learning/memory, and executive function and learning/memory, respectively, compared to those between region-wise accuracy maps (similarity measures of 0.655, 0.806, and 0.672, respectively). Conversely, the relevance scores showed higher similarity with region-wise accuracy map of the same domain compared to between-domain similarity measures (Fig. 3A). This suggests that relevance score-based analysis can capture fine-grained differences in FC-cognition association across cognitive domains and preserve the domain-specific FC-cognition associations.

**Fig. 3.**
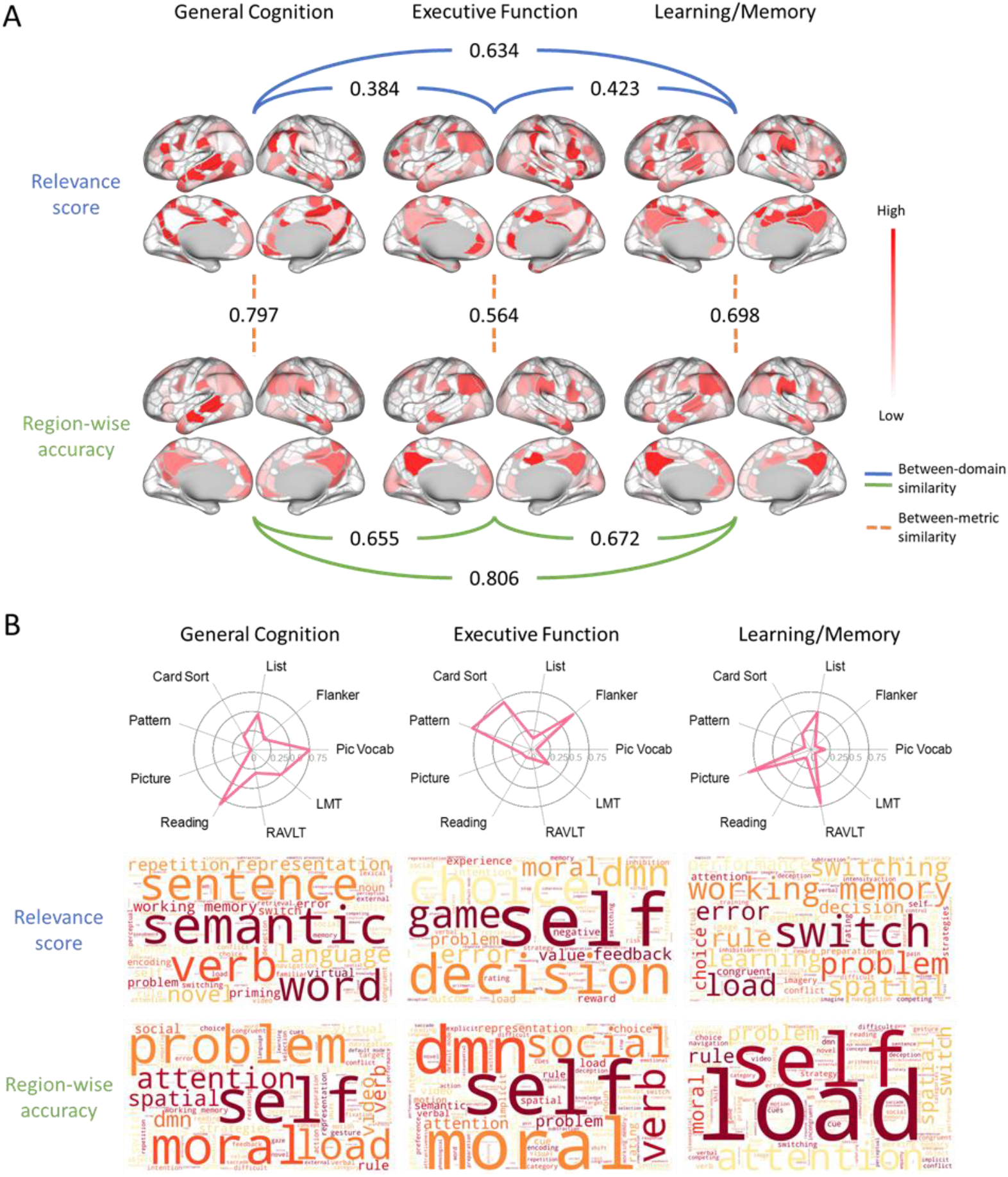
Comparison of FC-cognition associations quantified by different metrics. (A) FC-cognition association maps quantified using the interpretable predictive model’s relevance scores (top) and the region-wise ridge regression models’ prediction accuracy (bottom). For the latter, only brain regions with the top 50% prediction accuracy are shown to highlight regional differences. The number between the maps indicates their similarity. (B) Loadings of different cognitive assessments in the factor-analysis model across cognitive domains [30] (top). Meta-analytic functional decoding results for relevance score based (middle) and region-wise accuracy based (bottom) association maps across cognitive domains. Larger font size indicates stronger correlation with the specific topic.

To qualitatively interpret the differences in the FC-cognition associations identified by different metrics, we leveraged the meta-analytic functional decoding technique [39] to annotate each FC-cognition association map with its related functional topics. Specifically, an LDA-based meta-analysis combined with the NeuroQuery database [40] was utilized to link each FC-association map to its specific functional/behavioral topics. As illustrated in Fig. 3B, the topics highly correlated to the relevance scores for general cognition were “semantic”, “sentence”, and “word”, showing correspondence to the cognitive assessments with highest loadings for the general cognition (Reading, Pic Vocab, and LMT). The topics highly correlated to relevance scores for executive function included “self”, “decision”, and “choice”, highlighting the mental processes involved in executive functioning. Similarly, the topics highly correlated to relevance scores for learning/memory included “switch”, “working memory”, and “problem”, aligning with the target cognitive domain. These correlated topics exhibited both diversity across cognitive domains and sensitivity to the target domains. In contrast, the topics identified for the region-wise prediction accuracy included “self”, “problem”, “moral”, “dmn”, and “load”, exhibiting larger overlap across the cognitive domains compared to those identified for the relevance scores. These results further support that the relevance scores learned by the interpretable predictive model can capture fine-grained FC-cognition associations.

To validate the effectiveness of the learned relevance scores, we hypothesized that improved prediction performance could be achieved by integrating the standalone region-wise predictions (from the Region-avg model) with the relevance scores as weights. This weighted prediction is denoted as Relevance-informed prediction, and its prediction performance is demonstrated in Fig. 4. The correlation between measured and predicted values obtained by the Relevance-informed prediction was 0.451 ± 0.018, 0.163 ± 0.021, and 0.257 ± 0.023 for general cognition, executive function, and learning/memory, respectively. Improvements in prediction over the Region-avg model were found, especially for general cognition and learning/memory (*p*<0.04, corrected *t*-test), indicating that the relevance scores truly identified brain regions functionally linked to cognition. It is worth noting that the Relevance-informed prediction performed worse than the interpretable predictive model, underscoring the advantages of end-to-end learning for region-wise predictions and relevance scores.

**Fig. 4.**
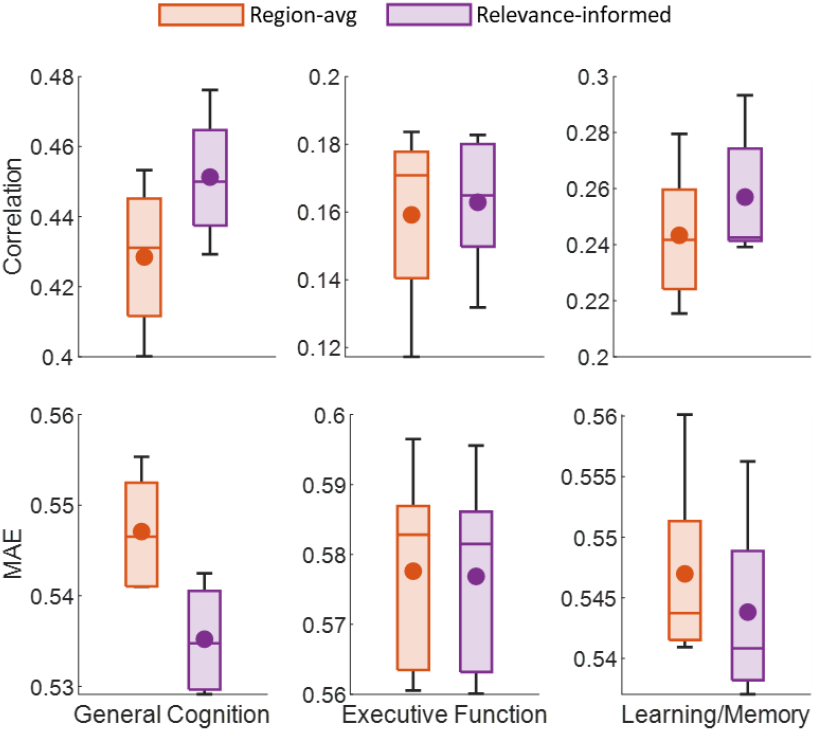
The relevance scores learned by the interpretable predictive model could improve the prediction accuracy for the region-wise prediction (Relevance-informed). The boxplot shows the distribution of prediction accuracy over five-fold cross-validation, with the ‘-’ and ‘•’ markers representing the median and mean, respectively.

### Generalization of relevance scores for longitudinal cognition prediction and external dataset validation

To further validate the functional meaningfulness of the relevance scores on a broader scale, we investigated whether the relevance scores could generalize and improve the performance of other cognitive prediction tasks. Two additional tasks were utilized. The first one was to predict the longitudinal cognitive measures at a 2-year follow-up using the baseline FC measures in ABCD cohort. The second task was to predict the cognitive measures from different domains using an external dataset from the HCP-D cohort. For each task, the baseline Region-avg model was trained and evaluated using five-fold cross-validation to obtain the standalone region-wise predictions for task-specific cognitive measures. The relevance scores learned from the ABCD baseline data were then utilized to integrate these predictions to generate Relevance-informed prediction results. It is worth noting that the interpretable predictive model was not re-trained for the task-specific cognitive measures, as the purpose of this validation was to evaluate the relevance scores’ generalizability to related cognitive prediction tasks.

For the longitudinal prediction tasks in the ABCD cohort, scores from seven cognitive assessments were evaluated (Reading, PicVocab, LMT, Pattern, Flanker, Picture, and RAVLT). When integrating the standalone region-wise predictions, each assessment was assigned to the cognitive domain with the maximal loading in the factor-analysis model [30], and the relevance scores for its corresponding domain were applied. Specifically, the relevance scores for general cognition were applied to the predictions of Reading, PicVocab and LMT, that for executive function were applied to Pattern and Flanker, and that for learning/memory were applied to Picture and RAVLT, respectively. For the prediction tasks in the HCP-D cohort, the averaged relevance scores for the general cognition, executive function, and learning/memory were applied to the cognitive scores to be predicted, including fluid cognition, crystalized cognition, and total cognition.

As shown in Fig. 5A, the relevance scores learned by the interpretable predictive model improved the prediction accuracy for all cognitive measures, especially for Reading, PicVocab, LMT, and Pattern (*p*<0.043, corrected *t*-test). The relevance scores also generalized well in predicting various cognitive scores from the HCP-D data. As shown in Fig. 5B, the Relevance-informed prediction significantly outperformed the baseline model Region-avg for predicting fluid cognition and total cognition (*p*<0.04, corrected *t*-test). These results further support that the relevance scores learned by the proposed model are robust and reproducible, capturing fine-grained FC patterns underlying cognition across multiple cognitive domains and cohorts.

**Fig. 5.**
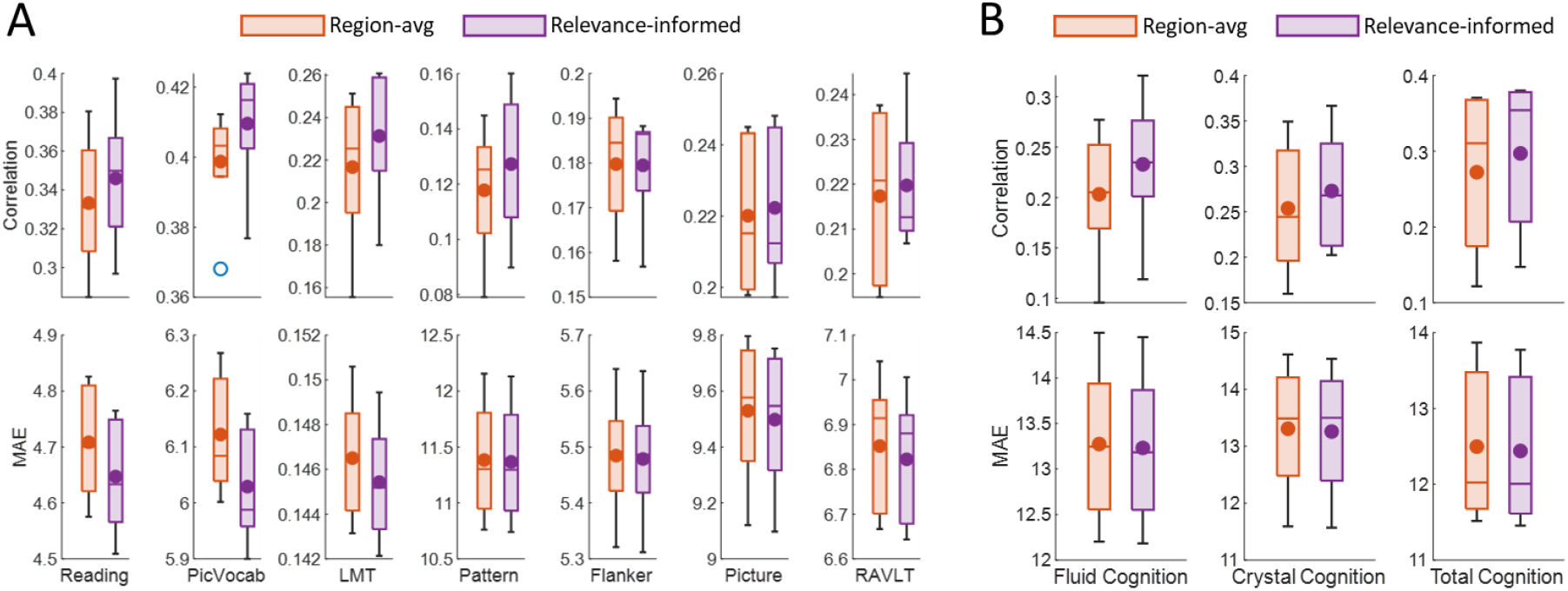
Relevance scores demonstrated strong generalization for both longitudinal cognition prediction and external dataset validation. (A) Predicting longitudinal cognitive measures (at 2-year follow-up) using baseline FC measures in the ABCD cohort. (B) Predicting cognitive measures in the HCP-D cohort. The boxplot shows the distribution of prediction accuracy over five-fold cross-validation, with the ‘-’ and ‘•’ markers representing the median and mean, respectively.

## Discussion

This study proposes an interpretable predictive modeling approach to identify fine-grained FC patterns predictive of behavioral traits at an individual level. By jointly capturing regional and whole-brain FC patterns and their associations with target behavioral traits in an interpretable predictive modeling framework, our model refines predictions and ensures consistency between region-specific and overall participant-level predictions and observed behavior measures. This interpretable modeling approach provides intrinsic model interpretability and enhances prediction accuracy, offering insights into the influence of brain FC on behavior. Applying this technique to a large sample of children, we identified informative, fine-grained FC-cognition association patterns. A quantitative comparison showed FC-cognition association patterns revealed by the relevance scores captured more domain-specific information than the region-wise accuracy. Moreover, relevance scores also improved longitudinal predictions of cognitive traits and cognitive predictions in an external cohort, demonstrating both robustness and biological relevance.

Predictive modeling is widely used to investigate connections between brain functional networks and cognition, with many studies employing machine learning algorithms on whole-brain FC measures as features to predict cognitive measures [5-9, 11]. While multivariate predictive modeling reveals significant FC-cognition associations, it is challenging to directly demonstrate the relative contribution of different brain regions for the prediction. Post-hoc analyses [10, 20, 41], such as summarizing the feature coefficients or feature importance at the region level or network level, are often utilized to highlight prominent brain regions and networks for the prediction. However, these post-hoc metrics may not accurately reflect the model’s predictive process. Region-level predictive models [12, 24] offer intuitive region importance evaluation via predictive performance, but fail to capture how multiple brain regions collectively influence behavior. A recent study utilizes stacking ensemble for brain age prediction based on brain structural imaging data by employing a two-step prediction strategy to integrate pre-trained regional predictions [42], but the whole-brain prediction may be limited by pre-trained regional predictions. Our end-to-end, interpretable modeling method addresses these limitations by using relevance score to integrate region-level predictions into an overall prediction, intrinsically explaining regional contributions. The relevance score informed prediction also improves performance over averaged region-level predictions. Furthermore, we observed that ensembles of region-level predictions generally outperformed whole-brain FC-based predictions, possibly due to the challenges high data dimensionality poses for the whole-brain models, limiting generalization.

Our learned relevance scores revealed that youth cognitive performance is primarily predicted by distributed brain regions across the association cortex, consistent with previous findings [5, 7, 21, 43, 44]. The relevant brain regions varied across cognitive domains. Specifically, the cingulo-parietal, retrosplenial-temporal, and dorsal attention networks were most predictive of general cognition. The cingulo-parietal and retrosplenial-temporal networks, along with the salience network, were most predictive of executive function, while the dorsal attention and cingulo-opercular networks were most predictive of learning/memory. A quantitative comparison of the FC-cognition associations, derived from relevance scores and region-wise accuracy, revealed that relevance scores capture more domain-specific information. All the results indicate that relevance scores provided additional information about FC-cognition associations and captured fine-grained differences across cognitive domains beyond region-wise prediction accuracy, accounting for the interplay across brain regions and their associations with cognition.

The learned relevance scores demonstrated robustness and generalizability. When used to integrate region-wise predictions from a baseline model, they improved prediction performance for both baseline and longitudinal cognitive measures in the ABCD cohort, and for relevant predictions in an external dataset across different cognitive domains. This supports the effectiveness of our model in identifying functionally relevant brain regions for youth cognition.

While our interpretable modeling shows promise in capturing prominent FC patterns associated with youth cognition, some limitations should be acknowledged. First, the linear model used for region-wise prediction may limit its ability to capture non-linear effects. A principled framework for selecting the optimal (linear or non-linear) predictive model for each region holds the potential to better characterize the FC-cognition associations. Second, while the identified relevance scores generalized well to the HCP-D cohort, the limited age range of the ABCD participants used for model training may not fully capture the developmental effects on cognition throughout childhood and adolescence. Additional investigation is needed to gain further insights into possible neurodevelopmental changes. Third, while this study focused on FC-cognition associations, integrating multi-modal imaging data and investigating the brain patterns predictive of cognition from a multi-dimensional perspective merit future investigation. Lastly, other than cognition, the proposed modeling framework can be extended to other behavioral traits and phenotypes. Future work will explore these applications.

In summary, we present a new strategy for interpretable predictive modeling of brain functional network and cognition. Through region-wise models and relevance scores, our method provides interpretable results at both regional and whole-brain levels, elucidating fine-grained FC patterns associated with youth cognition, while enhancing prediction accuracy and generalization.

## Acknowledgements

This study was supported by grants from the National Institute of Health: R01EB022573, R01AG066650, R01MH120482, R01MH113550, R01MH112847, R37MH125829, R0110078141. Additional support was provided by NIH U24NS130411, NIH RF1MH121867, the AE Foundation, the Center for Artificial Intelligence and Data Science for Integrated Diagnostics at Penn, and the Penn/CHOP Lifespan Brain Institute.

## Notes

### Competing Interest Statement

The authors have declared no competing interest.

